# The neighborhood of interaction in human crowds is neither metric nor topological, but visual

**DOI:** 10.1101/2022.08.18.504451

**Authors:** Trenton D. Wirth, Gregory C. Dachner, Kevin W. Rio, William H. Warren

## Abstract

Global patterns of collective motion in bird flocks, fish schools, and human crowds are thought to emerge from local interactions within a *neighborhood of interaction*, the zone in which an individual is influenced by their neighbors. Both *topological* and *metric* neighborhoods have been reported in birds, but this question has not been addressed in humans. With a topological neighborhood, an individual is influenced by a fixed number of nearest neighbors, regardless of their physical distance; whereas with a metric neighborhood, an individual is influenced by all neighbors within a fixed radius. We test these hypotheses experimentally with participants walking in real and virtual crowds, by manipulating the crowd’s density. Our results rule out a strictly topological neighborhood, are approximated by a metric neighborhood, but are best explained by a *visual* neighborhood with aspects of both. This finding has practical implications for modeling crowd behavior and understanding crowd disasters.

## Introduction

Large-scale patterns of coordinated motion are observed in many animal groups, including flocks of birds, schools of fish, herds of mammals, and crowds of humans^1–5^. It is widely believed that such global patterns of collective motion emerge from many local interactions between individuals in a process of self-organization^1,6,7^. Understanding collective motion thus depends on characterizing these local interactions^8,9^. First, what are the *rules of engagement* that govern how an individual interacts with a neighbor? Second, what is the *neighborhood of interaction* over which these rules operate and the influences of multiple neighbors are combined? Here we aim to characterize the neighborhood of interaction in human crowds.

Many mathematical models of collective motion assume rules of engagement based on hypothesized forces of attraction, repulsion, and velocity alignment^10–14^. Such models – including our own^15^ – typically average the influence of all neighbors within a *metric neighborhood* or zone of fixed radius (Figure 1A, shaded region), with neighbor influence often decreasing with metric distance^15–17^. In contrast, others have proposed a *topological neighborhood*^18–20^ (Figure 1A, dashed lines) in which an individual is influenced by a fixed number of nearest neighbors, regardless of their metric distance, and neighbor influence may decrease with ordinal number.

**Figure 1.**
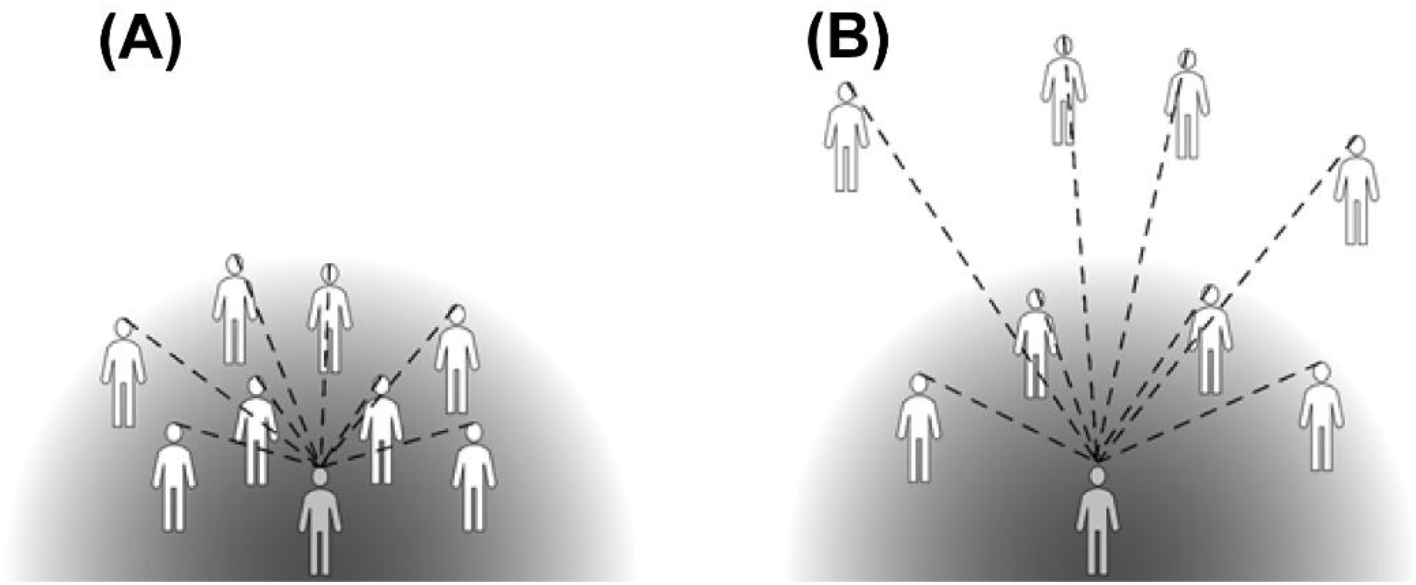
Testing the metric and topological hypotheses. (A) High density: Soft metric neighborhood (shaded gradient) predicts decreasing influence of neighbors (white figures) with metric distance from a pedestrian (gray figure), while a topological neighborhood (dashed lines) predicts decreasing influence with a neighbor’s ordinal distance. Metric and ordinal distance are correlated here. (B) Low density: The hypotheses are dissociated by manipulating crowd density. Whereas the soft metric neighborhood predicts that increasing neighbor distance will weaken their influence, the topological neighborhood predicts their influence will remain constant.

All of these models can be described as ‘omniscient’ because they assume a third-person view of the physical positions and velocities of all neighbors as input. We compare them with a new *visual neighborhood* model^21^ based on an embedded, first-person view in a crowd, which reflects both metric and topological distance^22^.

Evaluating these hypotheses is nontrivial, for metric distance (number of meters) and topological distance (ordinal number of neighbors) are naturally correlated. Yet the two hypotheses can be dissociated by varying group density. The metric hypothesis predicts that velocity alignment should depend on density, because the influence of neighbors increases with their physical proximity (Figure 1, shaded regions). In contrast, the topological hypothesis predicts that alignment should be independent of density, because neighbor influence only depends on ordinal number (Figure 1, dashed lines).

Observational studies of bird flocks have found empirical support for both hypotheses. Starlings appear to possess a topological neighborhood^2,18^, for the ordinal range of interaction remains constant at 6-7 neighbors despite natural fluctuations in flock density. In contrast, chimney swifts appear to have a metric neighborhood^23^, for the physical range of interaction is constant over variations in density (see also^24^). Specifically, alignment with neighbors is maximal at 1.4 m, independent of nearest-neighbor distance, and decreases with metric distance, whereas alignment with the *n*^th^ nearest neighbor depends on its metric distance.

To date, this question has not been answered in humans, for the existing data do not distinguish the hypotheses. The answer is of central importance for modeling crowd dynamics, simulating emergency evacuations, and understanding crowd disasters such as jams, crushes and stampedes^3,25–28^. We test the hypotheses by manipulating the density of virtual and real crowds, perturbing the heading (walking direction) of a subset of neighbors, and measuring the participant’s heading response. The metric hypothesis predicts that varying the density of neighbors will influence the heading response, whereas the topological hypothesis predicts that density will have no effect.

Specifically, we manipulate the distance of the perturbed and unperturbed neighbors in a virtual crowd so the metric hypothesis predicts a stronger (first experiment) or weaker (second experiment) heading response with a higher density. We find significant effects of density in the predicted directions. To generalize these results to real crowds (third experiment), we manipulate the density of human ‘swarms’ and analyze the degree of alignment. We find greater alignment in high-density swarms, whether plotted as a function of metric or topological distance.

These findings rule out a strictly topological neighborhood. The direction of the density effect is predicted by a metric neighborhood model^15^, but the quantitative results are best predicted by the visual model^21^ based on optical velocities and visual occlusion. We conclude that the neighborhood of interaction in humans is neither metric nor topological but visual, determined by the laws of optics.

## Results

We begin by describing models of metric, topological, and visual neighborhoods, then test them experimentally.

### Neighborhood models

#### Metric model

To describe a metric neighborhood, we used our empirical model of local interactions in human crowds^15^. The rules of engagement are derived from experiments on following in pairs of pedestrians, which found that the follower matches the heading direction and speed of the leader. The neighborhood of interaction is derived from experiments on a participant walking in a virtual crowd, which found that a pedestrian is influenced by a weighted average of neighbors (Equation 1a), where the weight decays exponentially with metric distance (Equation 1b):

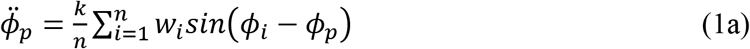

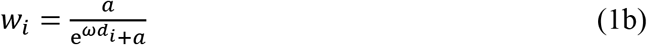

Specifically, a pedestrian *p*’s angular acceleration 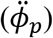 is proportional to the mean difference between *p*’s current heading (*ϕ_p_*) and that of each neighbor (*ϕ_i_*), where *n* is the number of neighbors within a 5m radius and a 180° field of view. The gain or stiffness parameter *k*=3.15 was fit to the pedestrian following data^29^. The weight of each neighbor *w_i_* decreases exponentially with metric distance *d_i_*, and the decay rate *ω*=1.3 and constant *a*=9.2 were fit to three trials of human ‘swarm’ data.

This results in a metric neighborhood with a ‘soft’ radius that asymptotes to zero around 4-5m, determining the range of interaction (Figure 1, shaded regions). According to the model, *p*’s heading direction stabilizes on the mean heading in the neighborhood. The physical proximity of neighbors determines the strength of attraction, and hence the turning rate and relaxation time of the heading response. An analogous equation for linear acceleration controls *p*’s speed^15^.

#### Topological model

A topological neighborhood is similarly based on a weighted average of neighbors (Equation 1a), but in this case the weight is a function of the topological distance of each neighbor (ordinal number *i*) rather than their metric distance:

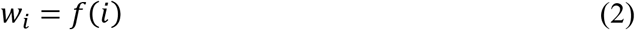

An integer number of neighbors, *i* = [1,…*,n*], is taken as input, where *n* determines the topological range of interaction. We do not try to estimate this ordinal function here, for the topological hypothesis can be tested qualitatively.

#### Visual model

The visual model^21^ is also based on a weighted average of neighbors, but it replaces omniscient variables (distance, heading, speed) with visual variables (angular velocity, rate of optical expansion, visibility):

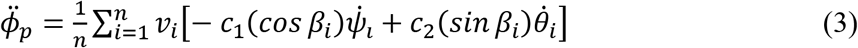

Specifically, pedestrian *p*’s heading is controlled by canceling the angular velocity 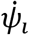 and expansion rate 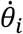 of all visible neighbors (*i* = 1… *n*). These two optical variables trade off as cosine and sine functions of the neighbor’s eccentricity *β_i_* in the field of view, which is centered on *p*’s heading direction. For example, if a neighbor directly ahead of *p* turns right, this generates a rightward angular velocity but little optical expansion; whereas if a neighbor on *p*’s left turns right, this generates an optical expansion but little angular velocity (see ref^21^ for details). The constants *c_1_* = 14.38 and *c_2_* = 59.71 were fit to data on pedestrian following^29,30^. A complementary equation controls *p*’s speed, based on the same optical variables^21^.

Critically, optical velocities 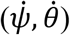 decrease with metric distance *d* as *tan*^-1^(1/*d*), in accordance with Euclid’s Law of visual angle, thus eliminating an explicit distance term (Equation 1b). In addition, nearer neighbors tend to visually occlude farther neighbors, depending on their visual directions, ordinal numbers, and metric separation in depth^22,31,32^. The model weights each neighbor in proportion to their *visibility*, which ranges from *v_i_* = 0 (fully occluded) to *v_i_* = 1 (fully visible); neighbors below a visibility threshold *v_t_* < 0.15 (fit to experimental data) are set to zero, and *n* is the number of neighbors at or above threshold. The neighborhood’s range of interaction is limited by the complete occlusion of farther neighbors, which varies with density and crowd configuration^22^.

The visual model thus depends on both metric and topological distance, but the neighborhood of interaction is determined by the laws of optics. The model stabilizes on the mean heading in the visual neighborhood, and the attraction strength, turning rate, and relaxation time are determined by the visibility of neighbors and the magnitudes of their optical motions.

### Experiments in Virtual Crowds

We tested the neighborhood hypotheses by varying the density of a virtual crowd. This allowed us to manipulate the behavior of virtual neighbors and measure their influence on a participant’s walking trajectory. Participants walked freely in a 12m x 14m area while viewing a group of 12 virtual humans in a mobile virtual reality headset. We asked participants to walk with the crowd and treat them as if they were real people. During each trial, we perturbed the heading (walking direction) of a subset of neighbors, all to the left (−10°) or to the right (+10°), and recorded the participant’s heading direction (the “heading response”).

High and low density crowds were created by positioning virtual neighbors at prescribed initial distances from the participant, and then randomly jittering their positions (Figure 2). On each trial, the virtual crowd appeared with their backs to the participant (Figure 2A,B); after 1s, a verbal “begin” command was played and the crowd accelerated forward for 3s to a walking speed of 1.0 m/s; 2s later the subset was perturbed, and the display continued for another 8 seconds. The participant’s head position in the horizontal plane was recorded, filtered, and used to compute the time series of heading for each trial. The final heading on each trial was the mean value between 4s and 6s post-perturbation. A mean time series was computed for each participant in each condition for analysis.

**Figure 2.**
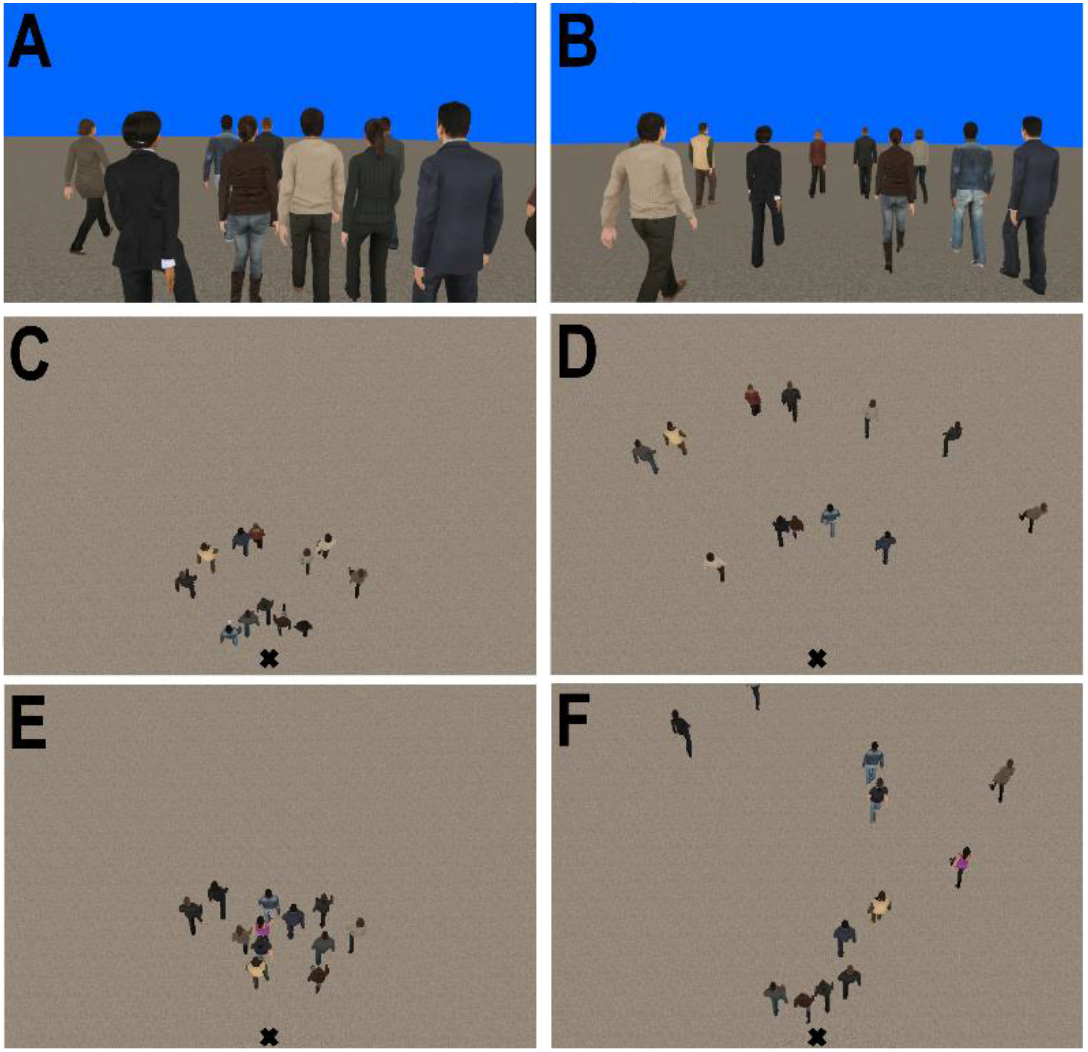
Virtual crowd displays. Participant’s view in the (A) High Density and (B) Low Density conditions of the first experiment. Bird’s-eye view in the (C) High Density and (D) Low Density conditions of the first experiment and (E, F) the second experiment. “X” indicates the participant’s position.

#### Heading responses increase with density when random neighbors are perturbed

In the first experiment, the heading of a random subset of the virtual neighbors (0, 3, 6, 9, or all 12) was perturbed on each trial. In the high density condition, five neighbors were initially positioned at 1.5m and seven at 3.5m (Figure 2C); in the low density condition, the initial distances were 3.5m and 7.5m (Figure 2D). (Speed was perturbed in another condition; see Supplementary Data 1 and Supplementary Figure 1).

According to the metric model (Equation 1), the participant is attracted to the mean heading in the neighborhood, which increases with the percentage of perturbed neighbors. Because nearer neighbors have higher weights, the model predicts that the attraction strength will be greater, the turning rate faster, and the relaxation time shorter at higher density. Consequently, the mean final heading after 4-6s should be larger in the high density condition than the low density condition, and this difference should increase with the percentage of perturbed neighbors (Figure 3A, dotted curves). In contrast, the topological hypothesis predicts no difference between the high and low density conditions.

**Figure 3.**
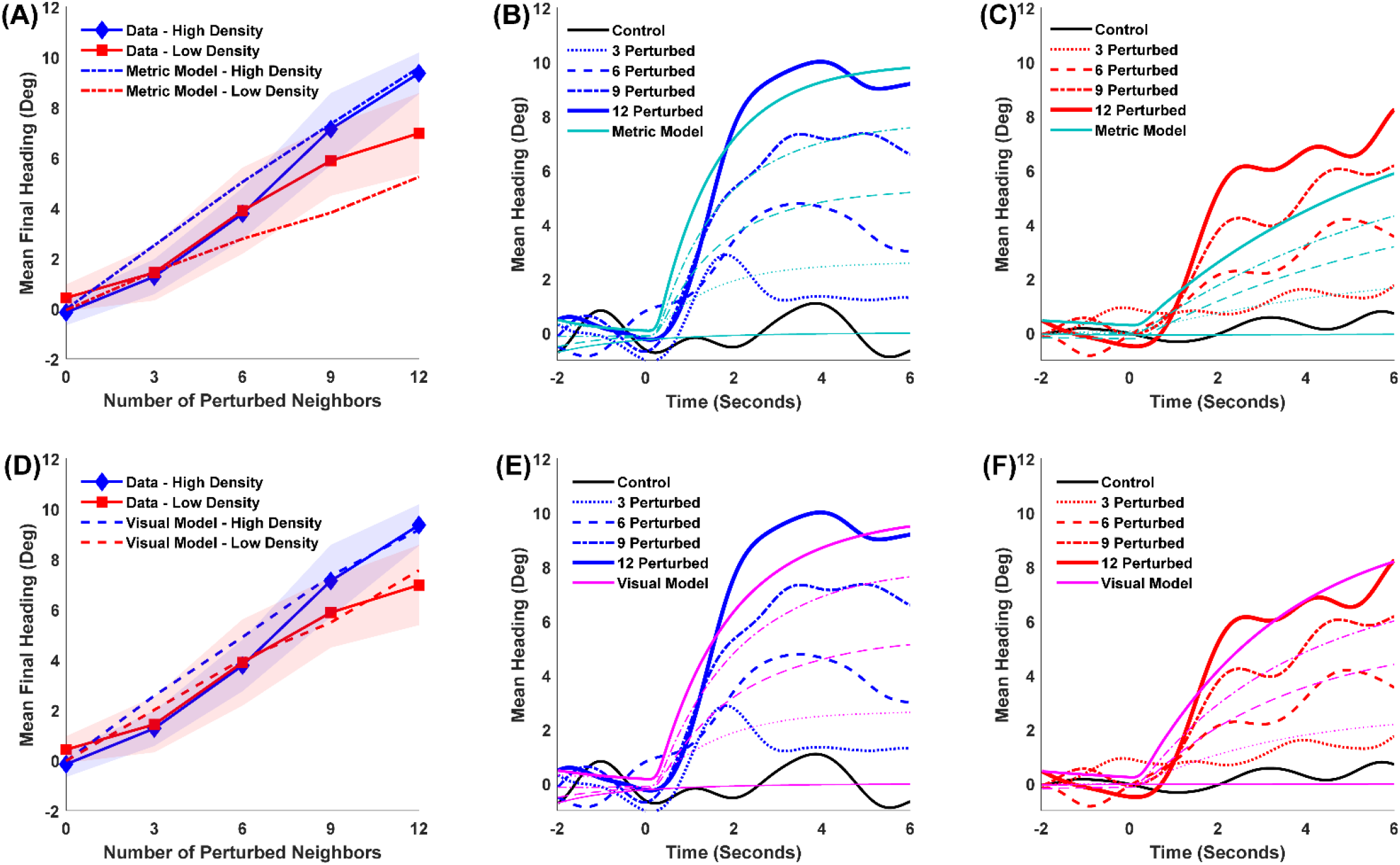
Results and simulations of the first experiment. (A) Mean Final Heading for humans and metric model as a function of the number of perturbed neighbors. Shaded regions represent the 95% confidence interval of the human data. (B) Mean time series of heading for human data and metric model in the High Density condition; curves represent the number of perturbed neighbors. (C) Same in the Low Density condition. (D,E,F) Same data with simulations of the visual model.

The results appear in Figure 3A (solid curves). As the number of perturbed neighbors increases, mean final heading becomes larger in the high density condition (blue) than the low density condition (red). A linear mixed effects (LME) regression analysis found that this interaction was significant: the effect of density increased with the number of perturbed neighbors (*χ*^2^(1) = 6.111, p = 0.0134) (see SI Methods for details on the statistical model). This significant dependence on density is contrary to the topological hypothesis. On the other hand, the metric model (dotted curves) is close to the human data, although it only lies within the 95% confidence intervals in four of the eight perturbation conditions, and undershoots the data in the low density condition (red).

Attraction strength is indicated by the time series of heading, where the post-perturbation slopes are steeper in the high density condition (Figure 3B, blue curves) than the low density condition (Figure 3C, red curves), increasingly so as more neighbors are perturbed. An LME regression showed that this three-way interaction (density x time x perturbed neighbors) is significant (*χ*^2^(1) = 4.163, p = 0.041), confirming that the turning rate is faster in the high density condition, as predicted by the metric model. In contrast, the significant dependence on density is inconsistent with the topological hypothesis.

##### Model simulations

To compare the metric model (Equation 1) with the human data quantitatively, we simulated each experimental trial with fixed parameters. The model agent was initialized with the participant’s position, heading, and speed 2s before the perturbation, the distance and heading of virtual neighbors were taken as input on each time step, and the agent’s heading time series was computed. We then calculated the agent’s mean time series in each condition, and compared it to the participant’s mean time series in the corresponding condition using the root of the mean squared error (RMSE).

Time series of heading for the metric model (cyan curves) are plotted together with the human data in Figure 3B (high density condition) and Figure 3C (low density condition). The model again appears to undershoot the data at low density. The mean RMSEm was 2.06° (leaving out the control condition).

We repeated these simulations using the visual model (Equation 3). In this case, the input to the model agent was the angular velocity, expansion rate, eccentricity, and visibility of each neighbor in the participant’s field of view, calculated from their position, heading, and speed at each time step. The model’s mean final heading (Figure 3D, dashed curves) is closer to the human data, particularly in the low density condition (red curves), and is within the 95% confidence interval of the data in six of the eight perturbation conditions. The mean time series for the visual model are plotted together with the human data in Figure 3E (high density) and Figure 3F (low density). The mean RMSE_v_ is 1.96°, closer to the human data than the metric model. To compare the relative strength of evidence for the two models, we computed Bayes Factors, yielding anecdotal evidence favoring the visual model overall (BF_vm_ = 1.42), with substantial evidence in the low density condition (BF_vm_ = 8.85). The visual model thus explains the human data as well as or better than the metric model.

Is this good model performance? Given that there is inherent noise in the data due to gait oscillations and measurement error, we estimated the limit on best performance by computing the RMSE between the participant mean time series in the control condition (0 perturbed neighbors) and a heading of 0°. This yielded a mean RMSE of 1.21°, indicating that the visual model is only 0.75° from the limit. Conversely, to estimate the worst performance for a model that does not respond to the input, we computed the RMSE between the participant mean time series in the perturbation conditions and a heading of 0°. This yielded a mean RMSE of 3.98°, indicating that visual model is much better than doing nothing. The visual model is thus near the high end of possible model performance, close to the human data.

##### Conclusion

The first experiment finds that participants have a stronger heading response in a higher density crowd. Specifically, when perturbed neighbors are in the majority and in closer proximity to the participant they exert a greater influence, producing a faster turning rate and a larger final heading. This significant density-dependence contradicts the topological hypothesis, which predicts that density should have no effect. The direction of the density effect is consistent with the metric hypothesis, but the data are better predicted by the visual model.

#### Heading responses decrease with density when nearest neighbors are perturbed

The first experiment found that the heading response increased with crowd density. But if the response depends on metric distance, we should be able to manipulate the proximity of unperturbed neighbors to elicit the opposite effect: a *decrease* in the heading response with higher density. The second experiment tested this prediction. Specifically, we held the metric distances of the four nearest neighbors constant and varied density by manipulating the distances of the other eight neighbors (Figure 2E,F). When the near neighbors are perturbed, the metric hypothesis predicts a weaker response in the high density condition than the low density condition. In contrast, the topological hypothesis predicts that the distance of the unperturbed neighbors should have no effect.

In this experiment, the heading of the nearest neighbors (0, 2, or 4) was always perturbed. The four nearest neighbors were positioned at fixed distances (1.5, 1.7, 1.9, 2.1 m) before jittering, while the remaining eight neighbors appeared at moderate distances in the high density condition (2.3 to 3.7 m, Figure 2E), and far distances in the low density condition (3.1 to 11.1 m, Figure 2F), and were never perturbed. The speed of the virtual crowd was increased slightly to a more comfortable walking speed (1.15 m/s), so the display continued for 5.4s post-perturbation and mean final heading was recorded between 2.4 and 4.4s. Otherwise, the procedure was the same as before.

The metric model predicts that the heading response should be reduced in the high density condition, because the unperturbed neighbors were closer and more influential. By contrast, in the low density condition the unperturbed neighbors were farther away and less influential, so the response to the perturbed neighbors should be stronger, yielding a faster turning rate and a larger final heading. In other words, the density effects should be the opposite of those observed in the first experiment.

The results for mean final heading appear in Figure 4A. It is clear that the density effect is reversed: final heading is now smaller the high density condition (solid blue curve) than in the low density (solid red curve). An LME regression confirmed a significant two-way interaction, such that the effect of density grows with the number of perturbed neighbors (*χ*^2^(1) = 5.54, p = 0.0186). This finding is similar to the first experiment but in the opposite direction, as expected by the metric hypothesis. On the other hand, a significant density effect contradicts the topological hypothesis. Yet the metric model (dotted curves) overshoots the data by a wide margin at both densities, lying outside the 95% confidence intervals in three of the four perturbation conditions.

**Figure 4.**
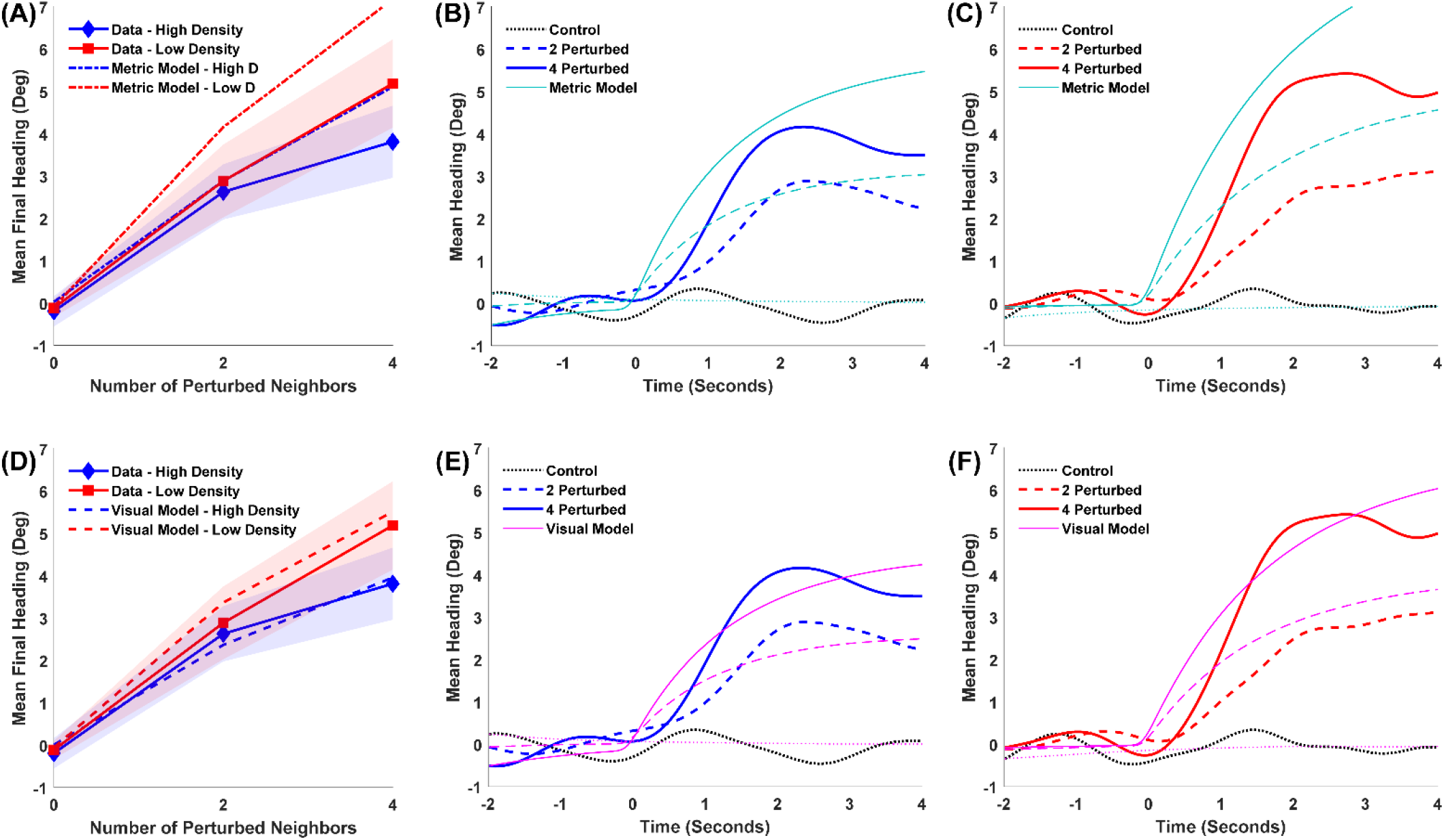
Results and simulations of the second experiment. (A) Mean Final Heading for humans and metric model as a function of the number of perturbed neighbors. Shaded regions represent the 95% confidence interval of the human data. (B) Mean time series of heading for human data and metric model in the High Density condition; curves represent the number of perturbed neighbors. (C) Same in the Low Density condition. (D,E,F) Same data with simulations of the visual model.

The effect of density on attraction strength is also reversed, for the slopes of the heading time series are shallower in the high density condition (Figure 4B, blue curves) than in the low density condition (Figure 4C, red curves). An LME regression found that the three-way interaction (density x time x number of perturbed neighbors) was significant (*χ*^2^(1) = 4.269, p = 0.0388), confirming a slower turning rate in the high density condition. In sum, the density effects were reversed, contrary to the topological hypothesis, but in the direction predicted by the metric hypothesis.

##### Model simulations

We simulated each trial with the metric model (Equation 1), as before. The mean time series of heading for the model (cyan curves) are plotted together with the human data in the high density (Figure 4B) and low density (Figure 4C) conditions. Leaving out the control condition, the mean RMSE was 1.48° (note the smaller error due to smaller turns in this experiment). Although the metric model generates the reversed density effect, it systematically overshoots the data.

Why might this be so? The metric model approximates the effect of distance with a fixed exponential decay term that was fit to a sample of human swarm data^15^. However, it does not take account of the actual optical velocities and visual occlusion in a particular crowd, and thus fails to generalize to other densities and distributions of neighbors. Because the visual model is predicated on these optical variables, it should generalize to the novel crowds in the second experiment.

We simulated the data with the visual model (Equation 3), as before. The model’s mean final heading appears in Figure 4D (dashed curves). Most importantly, it closely predicts the reverse density effect, falling within the 95% confidence interval for the data in all conditions. The mean time series of heading for the model are closer to the human data in both high density (Figure 4E) and low density (Figure 4F) conditions. Overall, the mean RMSE is 1.18° for the visual model, which is very strongly favored over the metric model (BFvb = 56.1). In addition, the performance of the visual model is only 0.43° from the inherent noise limit (mean RMSE = 0.75°), and it is better than doing nothing (mean RMSE = 2.55°). A visual neighborhood thus explains the human data better than a metric or topological neighborhood.

##### Conclusion

The second experiment found a significant density effect once again, but in the opposite direction of the first experiment. This density-dependence contradicts the topological hypothesis. The reversed density effect is consistent with the metric hypothesis, but the model overshoots the data in high and low density conditions. Both experiments are best explained by the visual model, which generalizes to new density and occlusion conditions because the neighborhood is based on optical variables rather than physical distance.

### Human ‘Swarm’ Experiment

To test whether our findings with virtual crowds extend to real crowds, a third experiment measured alignment in human ‘swarms’. Three groups of participants (N = 10, 16, 20) were instructed to walk about a large tracking area (14m x 20m), veering randomly left and right but staying together as a group, for 2-min trials. We manipulated the initial density of the group (high, low), and measured the difference in heading between pairs of participants to analyze alignment.

Each group participated in two trials at each density, for a total of 12 trials. Head positions in the horizontal plane were recorded with 16 motion-capture cameras, filtered, and used to compute the heading direction of each participant in each time step. This yielded approximately 11 minutes of usable data (frames in which all head positions were successfully recovered). We then measured the absolute heading difference (|Δ*ϕ_i,j_*|) and metric distance (*d_i,j_*) between pairs of participants *i* and *j* in each time step.

#### Alignment is greater in high density crowds

According to both the metric and topological hypotheses, the absolute heading difference between neighbors should increase with metric distance, because metric and topological distance are correlated. But the metric hypothesis predicts a smaller heading difference (greater alignment) in the high density condition, whether the data are plotted as a function of metric or topological distance. In contrast, the topological hypothesis predicts greater alignment in the *low* density condition when plotted as a function of metric distance, because the *n* nearest neighbors interact over longer distances. But any effect of density should disappear when the data are plotted as a function of topological distance.

We first checked that the density manipulation was successful by plotting a discrete probability density function for occupancy in each condition (Figure 5). A shift in color temperature between panels is apparent, indicating that a greater density was maintained in the high condition (Figure 5A, hot reds) than the low condition (Figure 5B, cooler oranges and yellows). The mean measured density (participants per square meter in every frame of data) was 2.1 ± 0.004 p/m^2^ (SEM) in the high condition and 1.72 ± 0.005 p/m^2^ (SEM) in the low condition. An LME regression analysis confirmed a significant effect of the high/low manipulation on measured density (*χ*^2^(1) = 2585.9, p < 0.001), although a significant interaction (high/low x time) indicated that the difference decreased over the course of a 2-min trial (*χ*^2^(1) =305.75, p < 0.001) (see Supplementary Figure 2).

**Figure 5.**
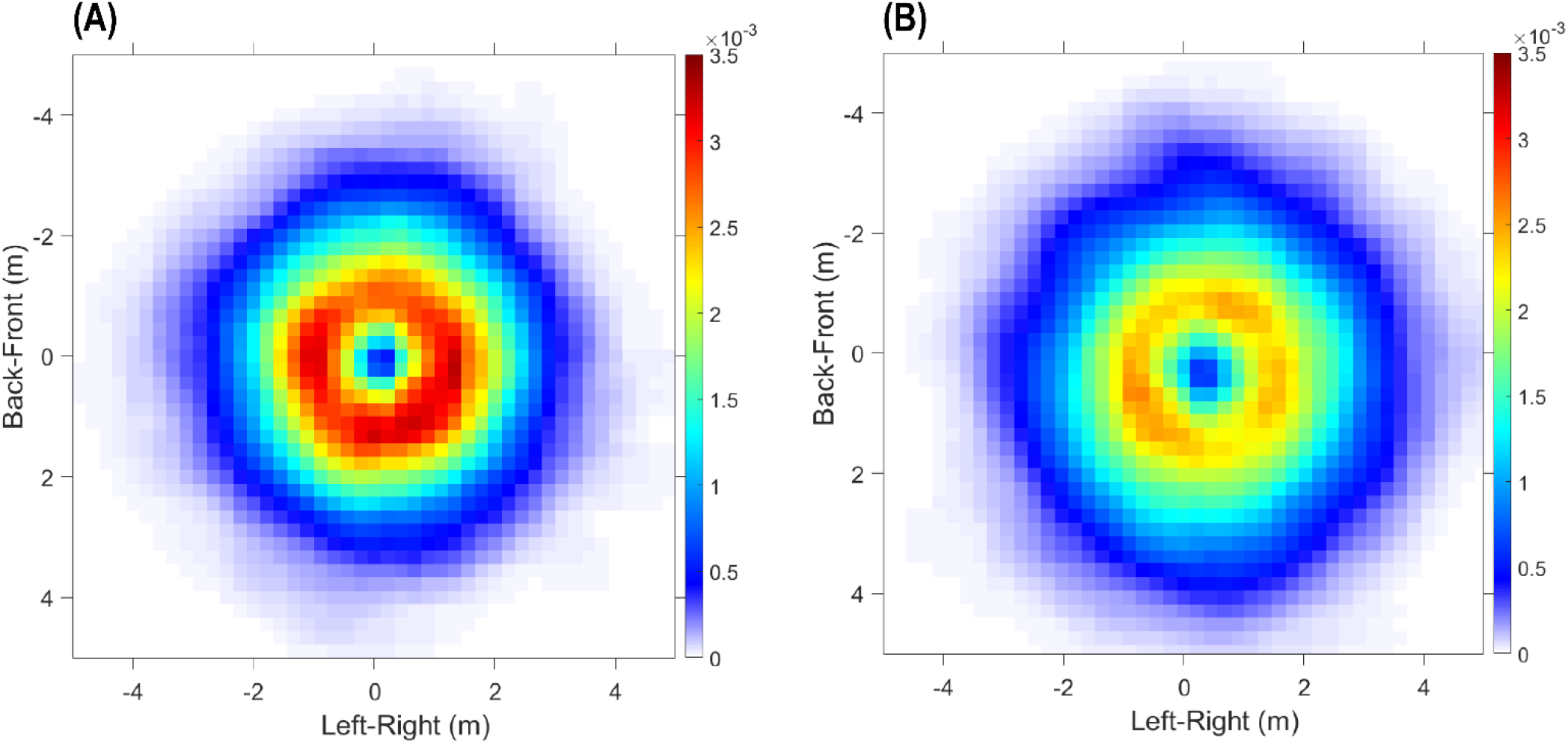
Occupancy PDFs for all human swarm trials, plotted relative to the focal participant nearest the centroid of the swarm. (A) High Density condition (mean 2.1 p/m^2^), 6 trials, 4.9 min of data. (B) Low Density condition (mean 1.7 p/m^2^), 6 trials, 6.2 min of data. Color temperature represents the discrete probability density of observing a participant in each 0.2m x 0.2m cell, with focal participant *p* at the origin, heading upward. Larger area of hot reds in A confirms the density manipulation.

To visualize the degree of alignment, we plotted heat maps of the mean absolute heading difference (|Δ*ϕ_i,p_*|) between the ‘focal’ participant *p* closest to the group centroid and each neighbor *i* (Figure 6). The metric hypothesis predicts greater alignment in the high density condition^32,33^, and indeed there is a larger region of cold blues (small heading differences) in the high density (Figure 6A) than the low density (Figure 6B) condition. On the topological hypothesis, one would expect the opposite, for the *n* nearest neighbors interact over longer distances in the low density condition.

**Figure 6.**
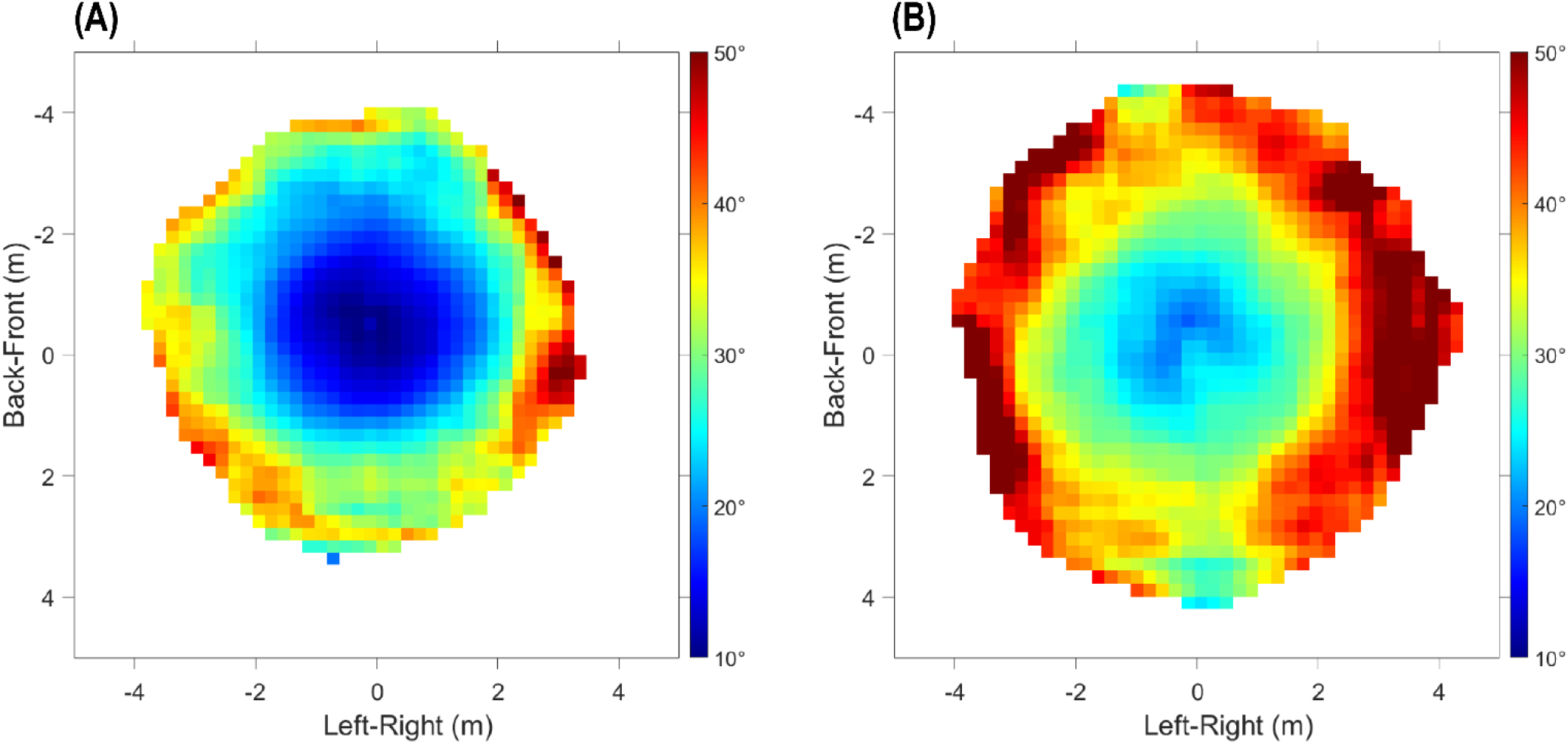
Heat maps of mean heading difference between the focal participant and each neighbor in human swarms. (A) High Density condition, 6 trials, 4.9 min of data. (B) Low Density condition, 6 trials, 6.2 min of data. Cells were only included in the heat map if they had at least 500 samples, or 8.33 seconds worth of data. Color temperature represents the mean absolute heading difference |Δ*ϕ_i,p_*| between the focal participant *p* nearest the swarm’s centroid (plotted at the origin, heading upward) and each neighbor *i* in the corresponding 0.2 x 0.2 m cell over all frames. Larger area of cold blues in A indicates greater alignment in the high density condition.

We then analyzed the dependence of alignment on metric distance. We computed the absolute heading difference between all pairs of participants *i* and *j* (|Δ*ϕ_i,j_*|), pruned extreme cases unlikely to interact, and then calculated the mean difference in consecutive 10s time intervals and 0.25m distance bins. An LME regression on heading difference in all trials confirmed a significant effect of metric distance (*χ*^2^(1) = 1508.1, p < 0.001); specifically, for every meter change in distance, there is a 4.74° ± 0.115° (SE) increase in the mean heading difference.

To test the neighborhood predictions, we sorted the heading differences (|Δ*ϕ_i,j_*|) by metric distance (0.25m bins) or by topological distance (ordinal number). When plotted as a function of metric distance (Figure 7A), overall the mean heading difference is smaller (greater alignment) in the high density condition (blue curve) than the low density condition (red curve). An LME regression on heading difference confirmed the density effect (*χ*^2^(1) = 7.35, p = 0.007), with a mean difference of 5.77° ± 1.36° (SE) between the high and low conditions. This finding is consistent with the metric hypothesis but contrary to the topological hypothesis. However, the interaction between density and distance was also significant (*χ*^2^(1) = 5.6, p = 0.018): the two curves cross at a distance of 2.75m, when the mean heading difference reaches 25°. At farther distances, the heading difference becomes larger at high density than low density, inconsistent with the metric hypothesis. This unexpected pattern is consistent with a visual neighborhood, for as distance increases there are more completely occluded neighbors in the high than the low density condition^22^ (see Supplementary Figure 3). Because a pedestrian is not influenced by these occluded neighbors, the mean heading difference become larger in the high density condition.

**Figure 7.**
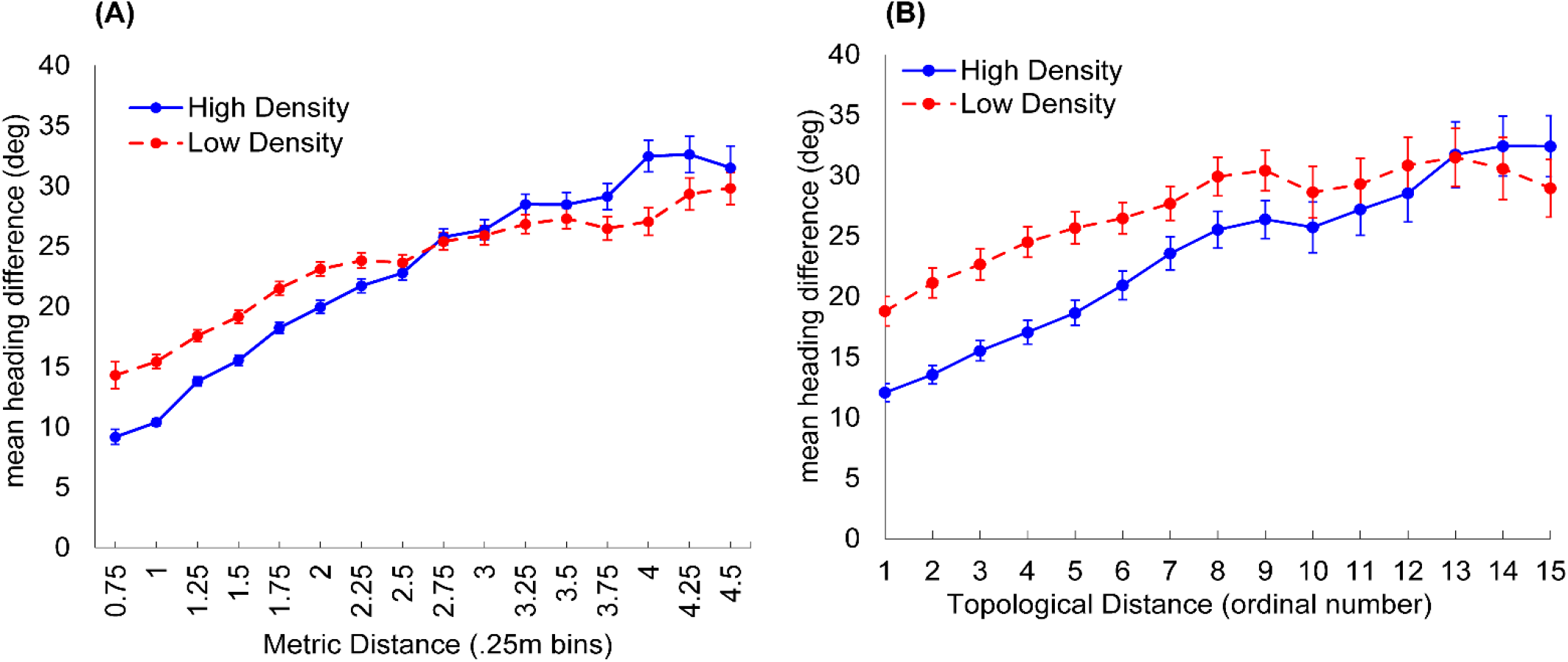
Mean absolute heading difference between all pairs of participants |Δ**ϕ_i,j_**| in human swarms, plotted as a function of (A) metric distance and (B) topological distance between *i,j*. The heading difference is smaller overall in the high density condition (blue) than in the low density (red) in both plots. Error bars represent the SEM, computed on the mean heading difference for all pairs of participants during each 10s interval in all trials (6 at each density). A. The high and low density curves represent all heading differences less than 50° for pairs less than 4.5m apart, yielding 4093 estimates in the high and 4022 in the low density condition. B. Data were re-sorted to obtain heading differences less than 50° for each of the 15 nearest neighbors in each 10s interval; metric distances ranged up to 8.3m. This yielded 549 estimates in the high and 528 in the low density condition.

When replotted as a function of topological distance (Figure 7B), the mean heading difference is still smaller in the high density (blue curve) than the low density (red curve) condition. This result indicates greater alignment of ordinally equidistant but physically closer neighbors, contradicting the topological hypothesis. An LME regression revealed that the density effect is significant (*χ*^2^(1) = 6.71, p = 0.010), although the interaction (density x ordinal number) is not, (*χ*^2^(1) = 1.67, p = 0.20).

##### Conclusion

The swarm experiment shows that heading alignment in real human crowds depends on density, whether plotted as a function of metric or topological distance. This finding provides decisive evidence against a topological neighborhood. The main density effect is consistent with a metric neighborhood, but the density x distance interaction supports a visual neighborhood.

## Discussion

Previous reports of collective motion in animal groups have found that some species, like starlings, possess a topological neighborhood that depends on ordinal distance, while others, like chimney swifts, have a metric neighborhood that depends on physical distance. The present research provides the first evidence that the neighborhood of interaction in human crowds is neither strictly topological nor strictly metric but visual, determined by the laws of optics.

The metric hypothesis predicts that varying density will affect the strength of interaction, because neighbor influence depends on metric distance. In contrast, the topological hypothesis predicts that varying density will have no effect, because neighbor influence only depends on ordinal number. The visual hypothesis predicts that responses will be influenced by density and visibility, factors that reflect both metric distance and ordinal number.

In three experiments we found that alignment reliably depended on density, specifically the proximity of perturbed and unperturbed neighbors. When random neighbors were perturbed, there was a stronger heading response at high density as the number of perturbed neighbors increased (also a stronger speed response). Conversely, when only the nearest neighbors were perturbed, there was a stronger heading response at *low* density, for unperturbed neighbors were farther away and exerted less influence. Measurements of human swarms also revealed a significant effect of density: we observed greater alignment at high density, regardless of whether the data were plotted as a function of metric or topological distance. The pattern of data thus qualitatively rules out a topological neighborhood, is in the expected direction for a metric neighborhood, but is more closely predicted by a visual neighborhood.

The visual neighborhood is determined by two factors, derived from the viewpoint of a pedestrian embedded in a crowd^21^. First, when a neighbor changes heading direction or speed, this generates corresponding optical motions in the pedestrian’s field of view. These optical velocities decrease with metric distance in accordance with Euclid’s law. Second, near neighbors tend to partially occlude far neighbors, such that visibility decreases with both ordinal number and metric separation in depth. The neighborhood’s range of interaction corresponds to the distance at which nearer neighbors completely occlude all farther neighbors, and thus varies dynamically with changes in density and visibility.

The visual model not only explains the density effects observed in the present experiments, it predicts the data in the second experiment much better than the metric model (Figure 4). Whereas the omniscient metric model describes the decay with distance using a fixed exponential function, the visual model explains this distance-dependence based on Euclid’s law and the geometry of occlusion. Because it is sensitive to variation in neighbor distance and visibility, the model generalizes to crowds with different densities and distributions of neighbors. The present results thus add support for a visual neighborhood in human crowds.

It is possible that previously observed topological and metric neighborhoods also have a visual basis. Notably, flocks of starlings and chimney swifts appear to have a different structure. Starlings^18^ maintain a spatial configuration by keeping a nearest neighbor in four visual directions in the field of view (±90° azimuth to the left and right, ±45° elevation up and down). The nearest neighbor in each quadrant would project the largest image and tend to occlude farther neighbors in that direction; with some positional drift, this would yield a topological neighborhood of 4-8 neighbors, consistent with the data. In contrast, roosting chimney swifts^23^ tend to align their velocities, and alignment decreases gradually with metric distance, from 1.4m to 4-5m. Heading responses in swifts might thus be governed by the same optical variables as in humans (Equation 3), which decrease with metric distance and greater occlusion. Thus, nominally ‘topological’ and ‘metric’ neighborhoods could be a consequence of the visual neighborhood of interaction.

We conclude that the neighborhood of interaction follows naturally from the laws of optics. The influence of visible neighbors decays with distance due to Euclid’s law, and the geometry of occlusion accounts for a further decrease in influence, until the range of interaction is limited by complete occlusion.

## Methods

### Virtual Crowd Experiments

#### Participants

Ten participants (5M, 5F) completed the first experiment, and 12 participants (7M, 5F) completed the second experiment; one additional participant was removed from the latter due to tracker error during data collection. All participants had normal or corrected-to-normal vision and none reported having a motor impairment. The research protocol was approved by Brown University’s Institutional Review Board, in accordance with the principles expressed in the Declaration of Helsinki. Informed consent was obtained from all participants, who were paid for their participation.

#### Equipment

The experiments were conducted in the Virtual Environment Navigation Lab (VENLab) at Brown University. Participants walked freely in a 12m x 14m tracking area, while viewing a virtual environment in a wireless stereoscopic head-mounted display (Oculus Rift DK1, Irvine CA; 90°H x 65°V field of view, 640 x 800 pixels per eye, 60 Hz refresh rate). Head position and orientation were recorded with a hybrid inertial/ultrasonic tracking system (IS-900, Intersense, Billerica MA), and used to update the display. The frame rate in the first experiment varied between 30-60 Hz, as did the tracker sampling rate; in the second experiment, the frame rate and sampling rate were constant at 60 Hz. The measured display latency varied between 50-67 ms.

#### Displays

The virtual environment consisted of a ground plane with a grayscale granite texture and a blue sky. A green start pole and a red orienting pole (radius 0.2m, height 3m) appeared 12.73 m apart (the start pole was reduced to 1.3m in the second experiment). The crowd consisted of 12 virtual humans (WorldViz Complete Characters) presented in the typical horizontal field of view (90°). In the first experiment only, 18 additional virtual humans were placed outside the field of view on two concentric circles to enhance the sense of immersion if the participant turned their head. The human models were animated with a walking gait with randomly varied phase. The racially diverse virtual crowd contained equal numbers of men and women.

In the first experiment, the 12 manipulated neighbors were initially positioned on two 90° arcs with the participant at the center, symmetric about the participant’s initial walking direction (toward the orienting pole). The arc radii were *r* =1.5m and 3.5m in the high density condition, or 3.5m and 7.5 min the low density condition. Five neighbors were placed at equal intervals on the near arc, and seven on the far arc. In the second experiment, the four nearest neighbors appeared at fixed initial distances (on arcs with *r* = 1.5, 1.7, 1.9, 2.1 m), and the nearest two or all four of them were perturbed. The other eight neighbors appeared on separate arcs spaced 0.2m apart in depth (*r* = 2.3, 2.5,… 3.7 m) in the high density condition, or 1m apart in depth (*r* = 3.1, 4.1,… 11.1) in the low density condition. The eccentricity *θ* of each neighbor was randomly selected from six equally spaced points on an 80° arc centered on the initial walking direction.

These initial positions were then jittered in polar coordinates, with the radial displacement Δ*r* randomly selected from a Gaussian distribution (*μ* = 0m, *σ* = 0.15m) and the angular displacement Δ*θ* from a separate Gaussian distribution (*μ* = 0°, *σ* = 8°). A different crowd configuration was generated for each trial; all participants received the same set of configurations, but virtual humans were randomly assigned to the positions.

During a trial, all virtual humans accelerated from a standstill (0 m/s) to a walking speed (1.0 m/s) on a straight path over a period of 3s following a sigmoidal function (cumulative normal, *μ* = 0, *σ* = 0.5s) fit to previous human data. After 2s, the heading direction of a subset of the 12 neighbors was perturbed by ±10°, all to the right or to the left, over a period of 0.5 s, following a similar sigmoidal function (*μ* = 0, *σ* = 0.083s). The display continued for another 8s. In the second experiment, crowd speed was increased to 1.15 m/s, closer to participants’ preferred walking speed, and the display thus continued for 5.4s.

The crucial manipulations were the following. In the first experiment, the perturbed subset (0, 3, 6, 9 or all 12 neighbors) was randomly selected from near and far neighbors. (The speed of the subset was similarly perturbed by ±0.3 m/s in a separate condition; see Supplementary Data 1). In the second experiment, only the nearest neighbors (0, 2, or 4) were perturbed. The four nearest neighbors always appeared at the same distances, and the density manipulation only affected the distances of the eight other neighbors.

#### Procedure

Participants were instructed to walk as naturally as possible, to treat the virtual pedestrians as if they were real people, and to stay together with the crowd. On each trial, the participant walked to the green start pole and faced the red orienting pole. After 2s, the poles disappeared and the virtual crowd appeared; 1s later, a verbal command (“Begin”) was played and the virtual crowd began walking. The display continued until the participant had walked about 12m (a duration of 12s in the first experiment and 10.4s in the second); a verbal command (‘End’) signaled the end of the trial. There were two practice trials to familiarize the participant with walking in the virtual environment. During this time, the participants could adjust the inter-ocular distance (IOD) of the HMD so that the display was clearly visible.

#### Design

*First experiment:* 5 perturbed subsets (0, 3, 6, 9, 12 neighbors) x 2 densities (high, low) x 2 perturbations (heading, speed). There were 8 trials per condition, for a total of 160 trials presented in a randomized order in two 1-hour sessions. The 80 heading-perturbation trials are reported in the text, and the results from the 80 speed-perturbation trials appear in Supplementary Figure 1. *Second experiment:* 3 perturbed subsets (0, 2, 4 nearest neighbors) x 2 densities (high, low). There were 16 heading-perturbation trials per condition, yielding a total of 96 trials presented in a randomized order in a 1-hour session.

#### Data Processing

For each trial, the time series of head position in the horizontal (X–Y) plane were filtered using a forward and backward fourth-order low-pass Butterworth filter to reduce the effects of oscillations due to the step cycle and occasional tracker error. Time series of heading direction and walking speed were then computed from the filtered position data. A 0.6 Hz cut-off was used for computing heading to reduce lateral oscillations on each stride, while a 1.0 Hz cutoff was used for computing speed to reduce anterior–posterior oscillations on each step. Right and left turn trials were collapsed by multiplying the heading angle on left turns by - 1. Speed-up and slow-down trials were collapsed by first (i) normalizing walking speed by subtracting the walking speed of unperturbed crowd (1 m/s) from participants’ speed time series, and then (ii) multiplying the normalized speed on slow-down trials by −1, to yield the absolute change in speed. Final heading and final speed were then computed as the average value during the last two seconds of each trial (4s to 6s post-perturbation in the first experiment, 2.4s to 4.4s in the second). To further mitigate the effect of gait oscillations, a mean time series was computed for each participant in each condition. Dependent measures included the mean final heading, and the mean time series of heading, for each participant in each condition (and the same for absolute speed change in the first experiment).

#### Statistical Analysis

We took a linear mixed effects (LME) regression approach, using the *fitlme* function (maximum likelihood approximation) in Matlab (R2019b). The dependent variable (e.g. heading) is regressed on predictor variables that may include categorical fixed effects (e.g. density), continuous fixed effects (e.g. time), and random effects (e.g. subjects, with unique intercepts). The residuals were inspected for any obvious heteroscedasticity or deviations from normality. Main effects and interactions were tested by comparing models in a step-down procedure that removes tested terms from the full model, using likelihood ratio chi-squared tests. Slopes are described for significant effects.

We performed two LME regression analyses: one on participant mean final heading, and the second on the participant mean heading time series (see Figure 3). Parallel analyses were performed on the speed data (see Figure S1).

### Human Swarm Experiment

#### Participants

One group of 10 participants, one group of 16 participants, and one group of 20 participants were tested in separate sessions as part of a larger study. The protocol was approved and informed consent was obtained as before, and participants were paid for their time.

#### Equipment

Head position was recorded in a large hall with a 16-camera infrared motion capture system (Qualisys Oqus, Buffalo Grove, IL) at 60Hz. The tracking area (14m x 20m) and starting boxes were marked on the floor with colored tape. Each participant wore a bicycle helmet with a unique constellation of five reflective markers on 30–40 cm stalks.

#### Procedure

Participants were instructed to walk about the tracking area at a normal speed, veering randomly left and right, while staying together as a group, for 2 min trials. Participants began each trial in shuffled positions in one of the starting boxes, corresponding to high and low density conditions: a 2×2m or 3×3m box for the 10-person group, a 3×3m or 4×4m box for the 16-person group, and a 4×4 or 7×7m box for the 20-person group. At a verbal ‘go’ signal, they began walking for 2 min, until a ‘stop’ signal. Each group received two trials in each density condition.

#### Design

3 groups (N=10, 16, 20) x 2 densities (high, low). There were 2 trials per condition, yielding a total of 12 trials with 24 min of raw data.

#### Data processing

The 3D position of the centroid of the markers on each helmet was reconstructed on each frame using a custom algorithm. Due to limits on the viewing volume and infrared reflections in the hall, there were many tracking errors, such that 100% of the helmets were recovered in 45% of all frames. The time series of head position in the 2D horizontal (*x,y*)plane was processed and filtered as before, and the heading direction of each helmet was computed on each time step in which it was successfully tracked; speed did not vary appreciably, and was not analyzed further.

We measured the density of the swarm in each frame as the number of participants per square meter (p/m^2^). Because this measurement depends on knowing the position of every participant, only frames in which 100% of the helmets were recovered were used in this analysis. We first determined the boundary of the (*x,y*) positions of the participants using Matlab’s *boundary* function, then computed the area of that polygon using the *polyarea* function, and finally calculated density by dividing the number of participants in that frame (*p*) by the area of the polygon (m^2^). Although this method overestimates absolute density somewhat, it captures the relative density in low and high conditions for each group (with constant N). For a robust estimate, we averaged the measured density of all frames in successive 10s segments on each trial, including only frames in which 100% of helmets were recovered. The mean density of the trials in each bin in the high and low conditions are plotted as a function of time bin in Supplementary Figure 2; the error bars represent the SE of the trial means in each time bin, where each bin includes 1-4 samples.

### Simulation Methods

Individual trials from the human swarm experiments were simulated in Matlab using the Runge-Kutta method (*ode45* function). The participant’s position, heading, and speed 2s before the perturbation were taken as the initial conditions. For the metric model (Equation 1), the input on each time step was the position, velocity, and speed of the virtual humans in the participant’s field of view in that trial. The speed of the virtual crowd was not perturbed in the second experiment, so the recorded time series of the participant’s walking speed on each trial was treated as input. For the visual model (Equation 3), the input was the angular velocity, optical expansion rate, eccentricity, and visibility of each virtual human, which were calculated from their positions on each time step. The output of both models were time series of the agent’s (*x,y*)position, heading, and speed for each trial.

#### Model comparisons

To compare the simulations with the human data, we first calculated the mean time series of heading (and speed) for each participant in each condition, and for the corresponding model agent. We then computed the mean absolute error (MAE) between each model agent and the participant time series in each condition. Finally, we compared the models to one another by calculating Bayes Factors (BFvm) based on the MAE between the model and each subject. Note that the variability in final heading is very small for the models because gait oscillations and tracker error were not simulated, so we compare the model means with 95% confidence intervals for the human data in the figures.

#### Model performance benchmarks

The performance of any model is limited by the inherent noise in the human data due to gait oscillations and tracker error. To benchmark the lower bound on error, we estimated the fluctuations in heading when walking on a straight path by computing the RMSE between each participant’s mean time series of heading on *control* trials (0 neighbors perturbed) and a heading of 0°. Conversely, to benchmark the upper bound on error – the failure of a model to respond to a perturbation – we estimated the error for a model that does not respond to the input by computing the RMSE between each participant’s mean heading time series on *perturbation* trials and a heading of 0°. These benchmarks indicate the range of model performance, from the best possible performance given the noise in the data to the performance of a model that does nothing. Of course, the performance of a model that responds inappropriately would be even worse.

## Data Availability

The experimental data that support the findings of this study are available in the Brown Digital Repository with the identifier “doi.org/10.26300/xk9b-5d55”^34^

## Acknowledgments

This research was supported by National Science Foundation (US) grants BCS-1431406 and BCS-1849446 to WHW, National Institutes of Health grants R01EY010923 and R01EY029745 to WHW, and T32 EY018080 to Brown University, and Link Foundation Fellowships to KWR, GDC, and TDW. Thanks to Adam Kiefer, Michael Fitzgerald, and the Sayles Swarm crew for help with crowd data collection; to Arturo Cardenas, Eugy Han and the student team for their help with data processing; and to Kei Yoshida for Supplementary Figure 3. Trenton Wirth is currently in the Biology Department at Northeastern University; Greg Dachner is currently with Uncommon Schools, New York; Kevin Rio is currently with Meta Reality Labs.

## Author Contributions

T.W., K.R., and W.W. designed research; T.W. and K.R. performed research; T.W. and G.D. processed and simulated the data; T.W. analyzed the data; and T.W. and W.W. wrote the paper.

## References

1 Couzin, I. D. & Krause, J. Self-organization and collective behavior in vertebrates. Advances in the Study of Behavior 32, 1–75 (2003).

2 Ballerini, M. et al. Empirical investigation of starling flocks: a benchmark study in collective animal behaviour. Animal behaviour 76, 201–215 (2008).

3 Helbing, D., Buzna, L., Johansson, A. & Werner, T. Self-organized pedestrian crowd dynamics: Experiments, simulations, and design solutions. Transportation Science 39, 1–24 (2005).

4 Parrish, J. K., Viscido, S. V. & Grunbaum, D. Self-organized fish schools: an examination of emergent properties. The Biological Bulletin 202, 296–305 (2002).

5 Sumpter, D. J. T. Collective animal behavior. (Princeton University Press, 2010).

6 Wirth, T. D. & Warren, W. H. Robust weighted averaging accounts for recruitment into collective motion in human crowds. Frontiers in Applied Mathematics and Statistics: Dynamical Systems 7, 761445 (2021). https://doi.org:10.3389/fams.2021.761445

7 Vicsek, T. & Zafeiris, A. Collective motion. Physics Reports 517, 71–140 (2012).

8 Gautrais, J. et al. Deciphering interactions in moving animal groups. PLoS Comput Biology 8, e1002678 (2012).

9 Sumpter, D. J. T., Mann, R. P. & Perna, A. The modelling cycle for collective animal behaviour. Interface Focus 2, 764–773 (2012).

10 Couzin, I. D., Krause, J., James, R., Ruxton, G. D. & Franks, N. R. Collective memory and spatial sorting in animal groups. Journal of Theoretical Biology 218, 1–11 (2002).

11 Czirók, A. & Vicsek, T. Collective behavior of interacting self-propelled particles. Physica A 281, 17–29 (2000).

12 Huth, A. & Wissel, C. The simulation of the movement of fish schools. Journal of Theoretical Biology 156, 365–385 (1992).

13 Reynolds, C. W. Flocks, herds, and schools: a distributed behavioral model. Computer Graphics 21, 25–34 (1987).

14 Helbing, D. & Molnár, P. Social force model of pedestrian dynamics. Physical Review E 51, 4282–4286 (1995).

15 Rio, K. W., Dachner, G. C. & Warren, W. H. Local interactions underlying collective motion in human crowds. Proceedings of the Royal Society B 285, 20180611 (2018).

16 Grégoire, G., Chaté, H. & Tu, Y. Moving and staying together without a leader. Physica D: Nonlinear Phenomena 181, 157–170 (2003).

17 Cucker, F. & Smale, S. Emergent behavior in flocks. IEEE Transactions on automatic control 52, 852–862 (2007).

18 Ballerini, M. et al. Interaction ruling animal collective behavior depends on topological rather than metric distance: Evidence from a field study. Proceedings of the National Academy of Sciences 105, 1232–1237 (2008).

19 Hemelrijk, C. K. & Hildenbrandt, H. Diffusion and topological neighbours in flocks of starlings: relating a model to empirical data. PLoS One 10, e0126913 (2015).

20 Hildenbrandt, H., Carere, C. & Hemelrijk, C. K. Self-organized aerial displays of thousands of starlings: A model. Behav. Ecol. 21, 1349–1359 (2010).

21 Dachner, G. C., Wirth, T. D., Richmond, E. & Warren, W. H. The visual coupling between neighbors explains local interactions underlying human ‘flocking’. Proceedings of the Royal Society B 289, 20212089. (2022).

22 Poel, W., Winklmayr, C. & Romanczuk, P. Spatial structure and information transfer in visual networks. Frontiers in Physics: Social Physics 9, 716576, 716571–716514 (2021).

23 Evangelista, D. J., Ray, D. D., Raja, S. K. & Hedrick, T. L. Three-dimensional trajectories and network analyses of group behaviour within chimney swift flocks during approaches to the roost. Proc. R. Soc. B 284, 20162602 (2017).

24 Lukeman, R., Li, Y.-X. & Edelstein-Keshet, L. Inferring individual rules from collective behavior. Proceedings of the National Academy of Sciences 107, 12576–12580 (2010).

25 Ngai, K. M., Burkle, F. M., Hsu, A. & Hsu, E. B. Human stampedes: a systematic review of historical and peer-reviewed sources. Disaster medicine and public health preparedness 3, 191–195 (2009).

26 Chraibi, M., Tordeux, A., Schadschneider, A. & Seyfried, A. in Encyclopedia of complexity and systems science (ed Robert A. Meyers) 1–22 (Springer Berlin Heidelberg, 2018).

27 Kinateder, M., Wirth, T. D. & Warren, W. H. in Crowd Dynamics, Volume 1: Theory, Models, and Safety Problems Modeling and simulation in science, engineering, and technology (eds N. Bellomo & L. Gibelli) 15–36 (Springer, 2019).

28 Kinateder, M. & Warren, W. H. Exit choice during evacuation is influenced by both the size and proportion of the egressing crowd. Physica A: Statistical Mechanics and its Applications 569, 125746 (2021).

29 Dachner, G. C. & Warren, W. H. Behavioral dynamics of heading alignment in pedestrian following. Transportation Research Procedia 2, 69–76 (2014).

30 Dachner, G. C. & Warren, W. H. A vision-based model for the joint control of speed and heading in pedestrian following. Journal of Vision 17, 716 (2017).

31 Pearce, D. J. G., Miller, A. M., Rowlands, G. & Turner, M. S. Role of projection in the control of bird flocks. Proceedings of the National Academy of Sciences 111, 10422–10426 (2014).

32 Strandburg-Peshkin, A. et al. Visual sensory networks and effective information transfer in animal groups. Current Biology 23, R709–R711 (2013).

33 Couzin, I. D., Krause, J., Franks, N. R. & Levin, S. A. Effective leadership and decision-making in animal groups on the move. Nature 433, 513 (2005).

34 Wirth, T., Dachner, G., Rio, K. & Warren, W. The neighborhood of interaction in human crowds is neither metric nor topological, but visual. Data sets. https://doi.org/10.26300/xk26309b-26305d26355 (2022).

