## Supplementary Information for "The neighborhood of interaction in human crowds is neither metric nor topological, but visual"

#### Supplementary Data 1

***Statistical analysis of the first experiment***

We performed two LME regression analyses: one on participant mean final heading, and the second on the participant mean heading time series (Figure 3). Parallel analyses were performed on the speed data (Supplementary Figure 1).

*Heading*. The first analysis tested the effect of the two-way interaction between density (categorical) and number of perturbed neighbors (continuous) on mean final heading, independent of time (Figure 3A, solid red and blue lines). The full model consisted of fixed effects for density, number of perturbed neighbors, and the two-way interaction term, and a fully specified random effect term with unique intercepts for each subject. Comparing the full model to a partial model without the two-way interaction of density and number of perturbed neighbors revealed that the interaction was significant ($\chi^2(1) = 6.111$, $p = 0.0134$). For each additional neighbor perturbed, mean final heading in the high density condition increased 0.245° ± 0.0848° (SE) more than in the low density condition. An effects structured list of model estimates appears in Supplementary Table 1A.

The second analysis tested the effect of the three-way interaction between density (categorical), number of perturbed neighbors (continuous), and time (continuous, centered on the perturbation), on heading (Figure 3B,C; red and blue curves, SI Table 1). The full model included the three single predictor variables, all two-way interaction terms, and the three-way interaction term as fixed effects, as well as a fully specified random effect structure providing unique intercepts for each subject. Comparing the full model to a model without the three-way interaction found the interaction to be statistically significant ($\chi^2(1) = 4.163$, $p = 0.041$). For each additional neighbor perturbed, the turning rate is 0.0393° ± 0.0172° (SE) per second faster in the high density than the low density condition. An effects structured list of model estimates appears in Supplementary Table 1B.

*Speed.* The influence of the random speed perturbation (0, 3, 6, 9 or 12 neighbors perturbed, at high and low densities) on the absolute change in walking speed is analogous to that of the heading perturbation described in the main body of the paper. We find, as predicted by Rio, et al.’s (2018) metric model, that the mean final change in speed is greater in the high than the low density condition (Figure S1 A). Using an LME regression, we tested the two way interaction between density (categorical) and number of perturbed neighbors (continuous) on mean final speed, with a fully specified random effects structure, and individual intercepts for each subject. The analysis reveals that the two-way interaction is significant, such that the final speed increased with the number of perturbed neighbors, moreso in the high density than the low density condition ($\chi^2(1) = 8.423$, $p = 0.00371$). For a full description of the fixed effects and the model, see Supplementary Table 1C.

Author Manuscript Draft

Moreover, the slopes of the time series (acceleration) are greater in the high density condition (Panel B) than the low density condition (Panel C), indicating a greater strength of attraction to the neighborhood speed. An LME analysis found that the three-way interaction (density x number of perturbed neighbors x time) is indeed significant ($\chi^2(1) = 11.353$, $p < 0.001$). This implies that the acceleration over time is greater in the high density than the low density condition, increasingly so as more neighbors are perturbed. These findings are consistent with a soft metric neighborhood, but contrary to a topological one. For a full description of the fixed effects and the model, see Supplementary Table 1D.

We performed simulations of the un-collapsed speed perturbation trials as described in the main text for heading simulations. The metric model (RMSE = 0.0401 m/s) and the visual model (RMSE = 0.0443 m/s) produce similar amounts of error, with anecdotal evidence favoring the metric model ($BF_{mv} = 1.229$). The similarity between the two models is likely due to the fact that visual occlusion does not vary much when the virtual neighbors change speed. We estimated the inherent noise due to gait oscillations by computing the RMSE between the participant mean time series in the control condition and a walking speed of 1m/s, the default walking speed of the crowd, yielding a mean RMSE = 0.0228 m/s. Finally, the "no response" estimate (a walking speed of 1 m/s in the perturbation conditions) yielded a mean RMSE = 0.0893 m/s; both models perform decisively better than doing nothing ($BF_{m0} > 100$, $BF_{v0} > 100$).

**Supplementary Data 2**

***Statistical analysis of the second experiment***

We performed LME regression analyses on the mean final heading and the participant time series of heading in the second experiment (Figure 4), using statistical models with the same structure as the first experiment.

*Heading*. For mean final heading, the two-way interaction between density and number of perturbed neighbors was significant ($\chi^2(1) = 5.54$, $p = 0.0186$). For each additional neighbor perturbed, mean final heading increased 0.324° ± 0.125° (SE) more in the high than the low density condition. An effects structured list of model estimates appears in Supplementary Table 2A.

For the time series of heading, the three-way interaction between density, number of perturbed neighbors, and time was significant ($\chi^2(1) = 10.158$, $p = 0.00144$). Critically, however, the direction of the density effect was reversed, as can be seen in Figure 4A,B: for each additional perturbed neighbor, the turning rate was 0.0645° ± .0285° (SE) per second faster in the *low* density condition than the high density condition. An effects structured list of model estimates appears in Supplementary Table 2B.

**Supplementary Data 3**

***Statistical analysis of the human swarms***

Author Manuscript Draft

To check the success of our density manipulation, we performed an LME regression analysis on the measured density in each frame (see Supplementary Figure 2). The full model included fixed effects for high/low condition, time (frame), the two-way interaction, as well as a random effect for the trial number; by comparing it to models without each subsequent term, we found significant effects of all three variables. Measured density decreased by -0.724 ± .014 $p/m^2$ (SE) between the high and low density conditions ($\chi^2(1) = 2585.9$, $p < 0.001$). This finding allowed us to treat density as a categorical variable in subsequent analyses. The mean measured density decreased over time by -0.0023 ± .00013 $p/m^2$ (SE) per second ($\chi^2(1) = 305.75$, $p < 0.001$), yielding a decrease in average density of -0.27 $p/m^2$ during a two-minute trial. Finally, there was also an interaction between density condition and time, such that the difference between the high and low conditions decreased by -0.005 ± .00013 $p/m^2$ (SE) per second ($\chi^2(1) = 756.57$, $p < 0.001$). Taken together, this implies that the high density condition dispersed by 0.6 $p/m^2$ over a two-minute trial. Although this finding suggests that a low density may be weakly preferred, it could also be a consequence of diffusion during swarming.

*Heading alignment.* To analyze alignment, we computed the absolute difference in heading between every recovered pair of participants ($|\Delta\phi_{i,j}|$) in every frame, as well as the distance between them. To reduce error in the data, we first removed extreme cases in which the pair had a mean heading difference greater than 50° (18.6%), indicating they were not interacting (cf. Figure 6), and outliers at distances greater than 4.5m (an additional 2.8%) (cf. the periphery in Fig. 5). For a robust estimate, we averaged the heading differences within successive 10s time bins and 0.25m distance bins, over all trials.

First, to demonstrate the relationship between distance between neighbors and their heading alignment, we performed an LME regression analysis on mean absolute heading difference with distance bin as a continuous fixed effect, and a random effect structure with participant pair and crowd size as correlated random intercepts, and time bin as an uncorrelated intercept. We found that for every meter increase in distance there was a 4.74° ± 0.115° (SE) increase in mean heading difference. We compared this model to a null model with the same random effect structure and found that the distance effect was significant ($\chi^2(1) = 1508.1$, $p < 0.001$).

Then, to test neighborhood predictions, we computed the mean absolute heading difference ($|\Delta\phi_{i,j}|$) when the data were sorted by metric distance (0.25m bins) or by topological distance (ordinal number) (Figure 7). We performed an LME regression on heading difference with metric distance bin, density condition, and their interaction as fixed effects, and a fully specified random effect structure including a random intercept for crowd size, and a correlated intercept for trial (a list of model estimates appears in Supplementary Table 3A). The heading difference decreases by 5.77° ± 1.36° (SE) from low to high density, with significantly greater alignment in the high density condition ($\chi^2(1) = 7.35$, $p = 0.007$). There is also a significant interaction between density and distance ($\chi^2(1) = 5.6$, $p = 0.018$).

We then re-sorted the data by topological distance and plotted the mean heading difference as a function of ordinal number (Figure 7B). Necessarily, there were fewer estimates in this case, for there was only one nth nearest neighbor in each 10s interval per trial. To reduce error, we again removed all estimates with a heading difference > 50° (13.95%), as well as outliers with an ordinal number >15 (an additional 3.96%). A similar LME analysis found a significant decrease in heading difference from low to high density, ($\chi^2(1) = 6.71$, $p = 0.010$), again indicating stronger alignment at the higher density (Supplementary Table 3B). There is also a significant

Author Manuscript Draft

increase in heading difference with ordinal distance ($\chi^2(1) = 8.12$, $p = 0.00437$), due to its correlation with metric distance. In this case, there is no interaction between density and topological distance, ($\chi^2(1) = 1.67$, $p = 0.20$). The finding of an effect of density (and hence metric distance) provides decisive evidence against the topological hypothesis.

Author Manuscript Draft

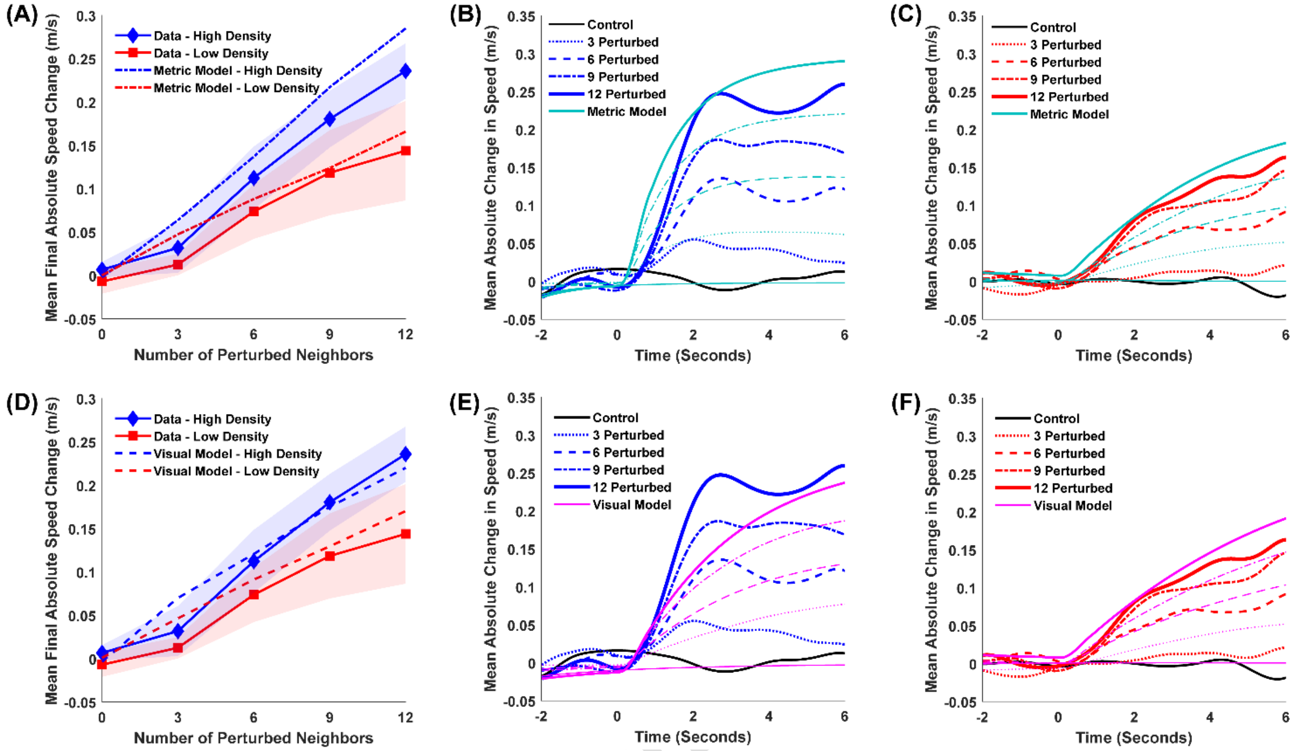


**Supplementary Figure 1.** Experiment 1: Results for walking speed with simulations of the metric and visual models. Panel A and D: Mean Final Absolute Speed Change (±0.3 m/s), with the metric model represented as the dash-dot line (A) and the visual model represented as the dashed line (D). Shaded regions represent the 95% confidence interval for the data. Panel B and C: Mean time series of absolute change in speed for each of the perturbation conditions, for high (B) and low (C) density, where the metric model is plotted in the cyan curves. Panel E and F: the same as panel B and C, except here the visual model is plotted in the magenta curves.

Author

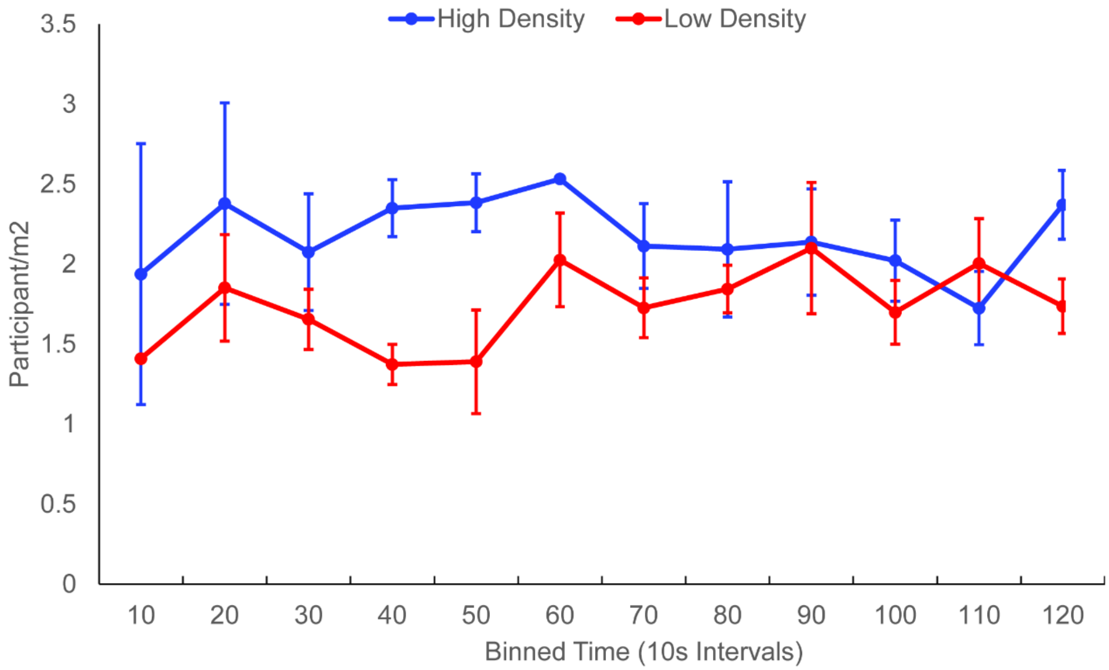


**Supplementary Figure 2.** Mean density (participants/$m^2$) as a function of time (10s intervals) in the human swarms (6 trials per density condition, 2 min each). Density was measured in each frame (60 Hz) and then averaged within successive 10s intervals for each trial. Error bars represent the SE of trial means in each time interval.

Author Ma

139

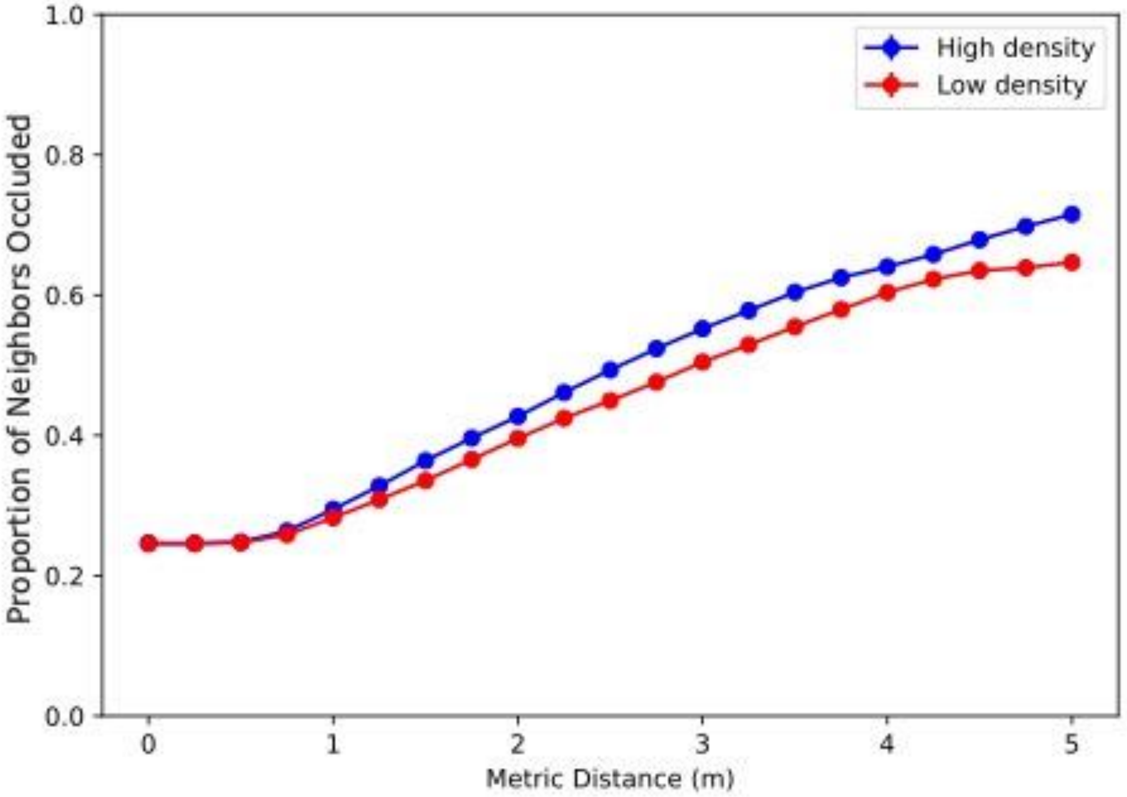


**Supplementary Figure 3.** Mean proportion of neighbors beyond each metric distance that are completely occluded ($v_i \leq 0.15$), in the high and low density conditions. Data based on all pairs of participants *i*,*j* in the human swarm, for neighbors *j* within *i*'s 180˚ field of view. [Thanks to Kei Yoshida for computing this figure.]

Author Manuscript

### Supplementary Table 1

*Experiment 1*

#### A) LME Regression: Mean Final Heading

*Formula*: Mean Final Heading~ Density*#Neighbors+(1+Density*Manip|Subject)

| Fixed Effects | Estimate | SE | t-statistic | p-value | 95% CI Lower | 95% CI Upper |
|---|---|---|---|---|---|---|
| Density | -0.457 | 0.266 | -1.720 | 0.089 | -0.984 | 0.070 |
| Number of Perturbed Neighbors (# Neighbors) | 0.707 | 0.061 | 11.582 | $p < 0.001$ | 0.586 | 0.828 |
| Density* #Neighbors | 0.123 | 0.042 | 2.889 | 0.005 | 0.038 | 0.207 |

#### B) LME Regression: Heading Time Series

*Formula*: Heading~ Density*#Neighbors*Time+(1+Density*#Neighbors*Time|Subject)

| Fixed Effects | Estimate | SE | t-statistic | p-value | 95% CI Lower | 95% CI Upper |
|---|---|---|---|---|---|---|
| Density | -0.074 | 0.155 | -0.478 | 0.632 | -0.378 | 0.230 |
| Number of Perturbed Neighbors | 0.128 | 0.022 | 5.865 | $p < 0.001$ | 0.085 | 0.171 |
| Time (Seconds) | -0.038 | 0.024 | -1.580 | 0.114 | -0.085 | 0.009 |
| Density* #Neighbors | 0.045 | 0.014 | 3.159 | 0.002 | 0.017 | 0.073 |
| Density*Time | -0.047 | 0.044 | -1.057 | 0.290 | -0.134 | 0.040 |
| #Neighbors*Time | 0.125 | 0.010 | 12.808 | $p < 0.001$ | 0.106 | 0.144 |
| Density* #Neighbors*Time | 0.020 | 0.009 | 2.272 | 0.023 | 0.003 | 0.037 |

#### C) LME Regression: Mean Absolute Change in Final Speed

*Formula*: Mean Final Speed~ Density*#Neighbors+(1+Density*Manip*|Subject)

| Fixed Effects | Estimate | SE | t-statistic | p-value | 95% CI Lower | 95% CI Upper |
|---|---|---|---|---|---|---|
| Density | 0.0026 | 0.0058 | 0.4440 | 0.6581 | -0.0090 | 0.0142 |
| Number of Perturbed Neighbors | 0.0169 | 0.0019 | 9.1070 | $p < 0.001$ | 0.0132 | 0.0206 |
| Density* #Neighbors | 0.0033 | 0.0010 | 3.4515 | $p < 0.001$ | 0.0014 | 0.0052 |

Author Manuscript Draft

**D) LME Regression: Speed Time Series**

*Formula*: Speed~ Density*#Neighbors*Time+ (1+Density*#Neighbors*Time|Subject)

| Fixed Effects | Estimate | SE | t-statistic | p-value | 95% CI Lower | 95% CI Upper |
|---|---|---|---|---|---|---|
| Density | 0.00521 | 0.00212 | 2.46250 | 0.01380 | 0.00106 | 0.00936 |
| Number of Perturbed Neighbors | 0.00271 | 0.00057 | 4.73620 | $p < 0.001$ | 0.00159 | 0.00383 |
| Time | -0.00245 | 0.00079 | -3.09240 | 0.00199 | -0.00401 | -0.00090 |
| Density* #Neighbors | 0.00060 | 0.00033 | 1.80780 | 0.07065 | -0.00005 | 0.00125 |
| Density*Time | -0.00084 | 0.00078 | -1.07160 | 0.28392 | -0.00238 | 0.00070 |
| #Neighbors*Time | 0.00311 | 0.00030 | 10.34800 | $p < 0.001$ | 0.00252 | 0.00370 |
| Density* #Neighbors*Time | 0.00082 | 0.00018 | 4.59590 | $p < 0.001$ | 0.00047 | 0.00117 |

Author Manuscript Draft

### Supplementary Table 2

*Experiment 2*

#### A) LME Regression: Mean Final Heading

*Formula*: Mean Final Heading~ Density*#Neighbors+(1+Density*Manip|Subject)

| Fixed Effects | Estimate | SE | t-statistic | p-value | 95% CI Lower | 95% CI Upper |
|---|---|---|---|---|---|---|
| Density | 0.039 | 0.146 | 0.265 | 0.792 | -0.252 | 0.330 |
| Number of Perturbed Neighbors | 1.162 | 0.106 | 11.009 | *p* < 0.001 | 0.952 | 1.373 |
| Density* #Neighbors | -0.162 | 0.062 | -2.592 | 0.012 | -0.287 | -0.037 |

#### B) LME Regression: Heading Time Series

*Formula*: Heading~ Density*#Neighbors*Time+(1+Density*#Neighbors*Time|Subject)

| Fixed Effects | Estimate | SE | t-statistic | p-value | 95% CI Lower | 95% CI Upper |
|---|---|---|---|---|---|---|
| Density | 0.029 | 0.083 | 0.351 | 0.726 | -0.134 | 0.192 |
| Number of Perturbed Neighbors | 0.292 | 0.038 | 7.647 | *p* < 0.001 | 0.217 | 0.367 |
| Time (Seconds) | 0.027 | 0.037 | 0.738 | 0.461 | -0.045 | 0.100 |
| Density* #Neighbors | -0.044 | 0.031 | -1.413 | 0.158 | -0.105 | 0.017 |
| Density*Time | 0.005 | 0.026 | 0.212 | 0.832 | -0.045 | 0.056 |
| #Neighbors*Time | 0.274 | 0.026 | 10.646 | *p* < 0.001 | 0.224 | 0.325 |
| Density* #Neighbors*Time | -0.032 | 0.014 | -2.264 | 0.024 | -0.060 | -0.004 |

144

Author Manuscript Draft

### Supplementary Table 3

*Experiment 3*

#### A) LME Regression: Heading Difference - Metric Distance

*Formula*: Heading Difference ~Binned Metric Distance*Density+(1+Binned M-Distance*Density|CrowdSize)+(1|Trial)

| Fixed Effects | Estimate | SE | t-statistic | p-value | 95% CI Lower | 95% CI Upper |
|---|---|---|---|---|---|---|
| Density | 5.776 | 1.359 | 4.249 | $p < 0.001$ | 3.111 | 8.440 |
| Binned Metric Distance | 9.150 | 0.928 | 9.865 | $p < 0.001$ | 7.332 | 10.968 |
| Density* Binned M-Distance | -2.216 | 0.528 | -4.198 | $p < 0.001$ | -3.250 | -1.181 |
| **Random Effects** | **Estimate (SD)** | | | | **95% CI Lower** | **95% CI Upper** |
| *Crowd Size* | | | | | | |
| Density | 0.389 | | | | 0.016 | 9.346 |
| Binned Metric Distance | 1.416 | | | | 0.626 | 3.201 |
| Density* Binned M-Distance | 0.780 | | | | 0.307 | 1.983 |
| *Trial* | | | | | | |
| Intercept | 2.091 | | | | 1.359 | 3.215 |

#### B) LME Regression: Heading Difference - Topological Distance

*Formula*: HdnDiff~Density*Topological Distance+(1+Density*T-Dist|CrowdSize)+(1|Trial)

| Fixed Effects | Estimate | SE | t-statistic | p-value | 95% CI Lower | 95% CI Upper |
|---|---|---|---|---|---|---|
| Density | -4.187 | 0.980 | -4.275 | $p < 0.001$ | -6.110 | -2.265 |
| Topological Distance | 1.475 | 0.322 | 4.584 | $p < 0.001$ | 0.843 | 2.106 |
| Density*T-Distance | 0.264 | 0.174 | 1.521 | 0.129 | -0.077 | 0.605 |
| **Random Effects** | **Estimate (SD)** | | | | **95% CI Lower** | **95% CI Upper** |
| *Crowd Size* | | | | | | |
| Density | 0.681 | | | | 0.054 | 8.550 |
| Topological Distance | 0.543 | | | | 0.256 | 1.151 |
| Density*T-Distance | 0.276 | | | | 0.120 | 0.632 |
| *Trial* | | | | | | |
| Intercept | 2.553 | | | | 1.604 | 4.066 |

Author Manuscript Draft
